# *anonymizeBAM*: Versatile anonymization of human sequence data for open data sharing

**DOI:** 10.1101/2021.01.11.426206

**Authors:** Christoph Ziegenhain, Rickard Sandberg

## Abstract

The risks associated with re-identification of human genetic data are severely limiting open data sharing in life sciences. Here, we developed *anonymizeBAM*, a versatile tool for the anonymization of genetic variant information present in sequence data. Applying *anonymizeBAM* to single-cell RNA-seq and ATAC-seq datasets confirmed the complete removal of donor-related genetic information. Therefore, the accurate generation of de-identified sequence data will re-enable open sharing in sequencing-based studies for improved transparency, reproducibility, and innovation.

## Main text

Modern omics methods have transformed life science research and especially sequencing-based approaches have gained in popularity^1,2^. Sharing sequencing data has been a major driver of innovation, and increased the pace of development in life sciences. Free access to raw sequencing data is essential for research transparency and for large-scale integration of published data. With the maturation of technologies like single-cell RNA sequencing^3^, many fields have progressed to broad applications in healthy and diseased primary human samples.

However, sequencing data generated from human donors comes with ethical concerns relating to data privacy. As sequencing data is re-identifiable given a large enough reference database^4^, it is paramount to protect the study individuals’ identity and genetic information, as reflected in recent legislation^5^. Therefore, authors are currently providing only the summarized data publicly, such as count tables in the case of RNA-seq, while the raw sequencing data are deposited in controlled-access repositories (e.g. dbGaP^6^ or EGA^7^). Alternative strategies, such as blockchain encryption algorithms have been proposed^8^, but have yet to reach widespread adoption^9^. While controlled-access repositories are important for protecting sensitive genomic data and sharing it only with researchers granted access, the heavily increased barriers in sharing sequencing data are severely limiting study transparency, reproducibility, innovation and development.

To lower the barriers in sharing sequence data, we propose, like others recently^10^, the deidentification of aligned reads by removing all information that reveals the identity and compromises the privacy of the donor (**Figure 1a**). Ideally, anonymized data can be openly shared and used in studies where the donor genetic information is not required. A data anonymization tool needs to correctly consider multiple types of genetic variation (SNPs, insertions, deletions, immune haplotypes, etc.), present in different sequencing-based data (e.g. RNA-seq, scRNA-Seq, DNA-Seq, ATAC-seq) from various sequencing platforms (Illumina, MGI, Pacific Biosciences, Oxford Nanopore Technologies) and strategies (e.g. single- or paired-end sequencing). At the same time, the remaining information (e.g. alignment positions, custom SAM tags) should be preserved as close to the original as possible, and the strategy should be agnostic to the sequence alignment tool used, for example STAR^11^ and BWA^12^ report alignments differently that affect the data anonymization process. In contrast to a recent study^10^, we reasoned it would be possible to write a single, versatile and intuitive tool for the anonymization of all kinds of genomics data, that can easily be integrated into existing infrastructures.

**Figure 1.**
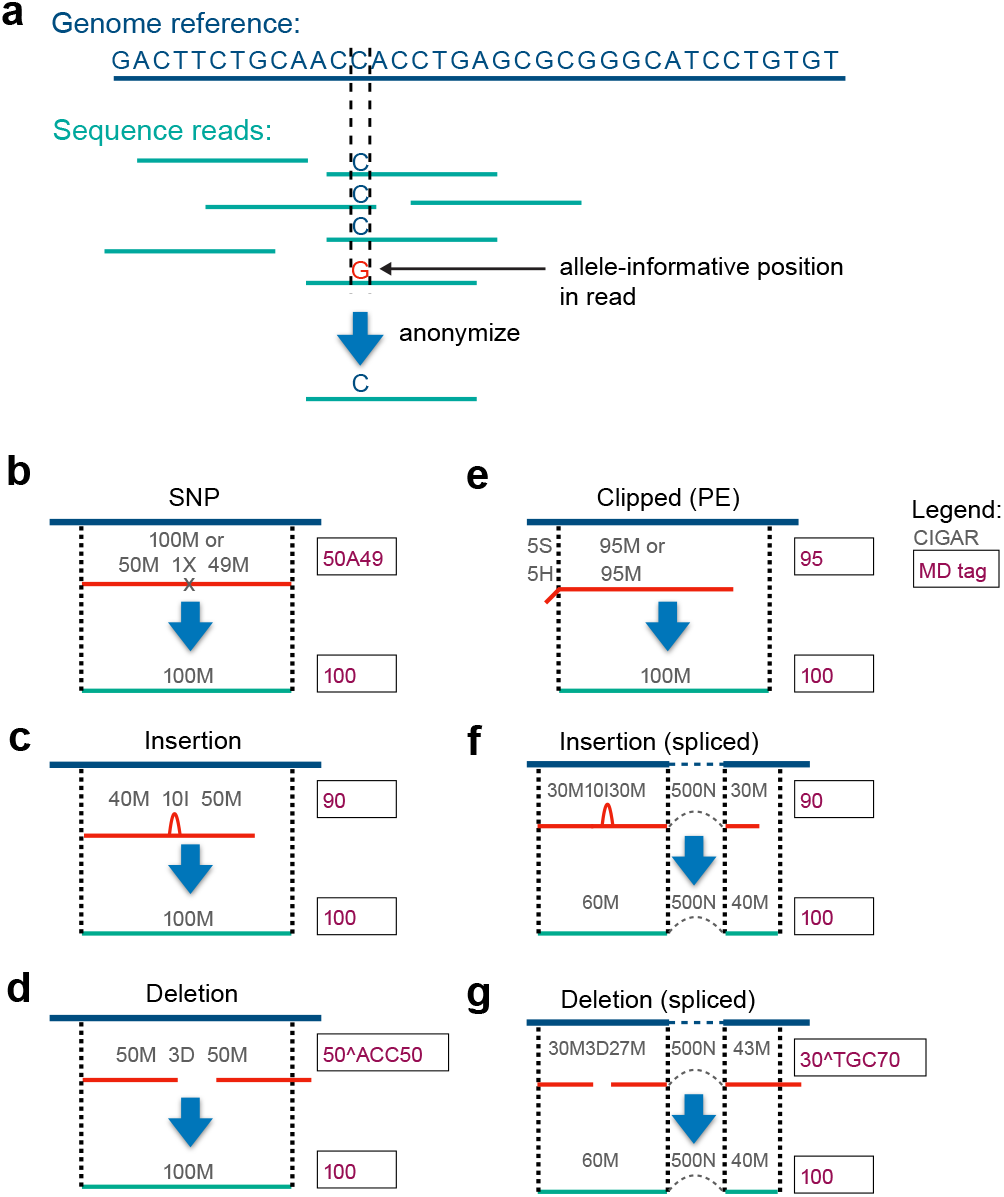
Schematic overview of the read anonymization procedure. (**a**) Sequenced reads typically harbor identifying genetic variants that are deviating from the reference genome. Data sanitation removes such variants by replacement with the reference sequence. (**b**) Schematic representation of an aligned 100 bp read (red) with alternative allele at SNP position is corrected to reference base in anonymized read (blue). Raw and corrected CIGAR strings (with operator descriptions below) are shown over reads, and raw and corrected MD tags are shown to the right. (**c**) Aligned 100 bp read (red) with donor-specific insertion (relative to reference), with information as in (b). (**d**) Aligned 100 bp read (red) with donor-specific deletion (relative to reference), with information as in (b). (**e**) Aligned 100bp read pair sequence (paired-end data) that was clipped at the 5’ end relative to reference. Note the correction in alignment end position as a result of the removal of clipped positions. (**f**) Spliced alignment that additionally contains an insertion not present in reference. The spliced alignment information is preserved during the sanitation of genetic variation. (**g**) Spliced alignment that additionally contains a deletion not present in reference. The spliced alignment information is preserved during the sanitation of genetic variation. Cigar string operators: M, alignment match; I, insertion; D, deletion; N, skipped in reference; S, soft clipping; H, hard clipping; P, padding; =, sequence match; X, sequence mismatch.

In brief, data anonymization can be achieved for sequenced and aligned reads stored in the ubiquitous BAM (Binary Sequencing Alignment Map) format by replacing donor genetic variation with the sequence of the reference genome (**Figure 1a**). Anonymization cannot be applied to the unmapped reads in the BAM file, which are therefore discarded. In the simplest case, a fully aligned sequence read containing a single nucleotide polymorphism (SNP) is deidentified by replacement with the reference base (**Figure 1b**). In the case of insertions in the donor sequence, the non-reference portion of the sequenced read is discarded and the deidentified sequence is extended by the length equal to the insertion while keeping the 5’ mapping position intact (**Figure 1c**). Conversely, deletions are anonymized by inserting the missing reference sequence and removing the equal numbers of bases from the 3’ end of the de-identified sequence read (**Figure 1d**). Portions of the read that may be soft- or hard-clipped are replaced by reference sequence, as the non-matching sequence could stem from both technical artifacts such as adapters but also from genetic variation. In the case of reads starting with clipping, for single-end sequencing data the reference position of the read start is adjusted, however this is not possible for paired-end reads because it would invalidate the mate-pair information (TLEN and PNEXT fields). Instead for paired-end reads, the clipped sequence portion is added to the end of the read (**Figure 1e**). In RNA-sequencing data, spliced alignments are common and reads can span several exon-exon junctions. For these reads, both the 5’ mapping position and the location of the splice junctions are preserved when sanitizing the sequence data of insertion and deletion events (**Figure 1f-g**). In addition to the actual sequence field itself within the BAM file, information on the presence of genetic variation in the donor individual is stored in the CIGAR string and accessory fields. Therefore, the CIGAR field and MD tag are consequently corrected while removing each type of genetic variation (**Figure 1**). Genetic information could be inferred from additional accessory fields such as alignment score, mapping quality and other alignment information (e.g. number of hits NH). To solve this, we update or remove these fields while leaving other auxiliary tags (e.g. sample information, cell barcode, gene identity, or custom flags) in place. While *anonymizeBAM* aims to report data as close to the input sequence data as possible, resolving indel variation results in out of phase quality scores with respect to the base calls. The resulting anonymized BAM file is fully compliant with the Sequence Alignment Map (SAM) specifications^13^ (as confirmed by picard-tools validation; see methods) and thus smoothly compatible with existing bioinformatics tools and pipelines. We implemented the anonymization strategy in an open-source stand-alone Python script, called *anonymizeBAM*, that only requires the input BAM file and reference sequence (i.e. the genome or transcriptome reference used for the alignment).

To illustrate the effectiveness of our program, we analyzed a recently published dataset of single-cell RNA sequencing (scRNA-seq) data generated from five equally abundant cell lines^14^ via the popular 10x Genomics^15^ protocol. After preprocessing the raw data with zUMIs^16^, we summarized SNP coverage and assigned the 2,937 high-quality (>= 66% exonic, >= 25,000 reads) cells to their donor of origin using cellsnp-lite^17^ and vireo^18^. Clearly, each of the cells in the main clusters could be assigned to a single donor corresponding to one of the cell lines (**Figure 2a**). Next, we processed the BAM file with *anonymizeBAM* and repeated the donor assignment using the same settings, which resulted in the inabilityto assign any donor structure (**Figure 2b**) due to the complete lack of donor-specific allele information (**Figure 2c**).

**Figure 2.**
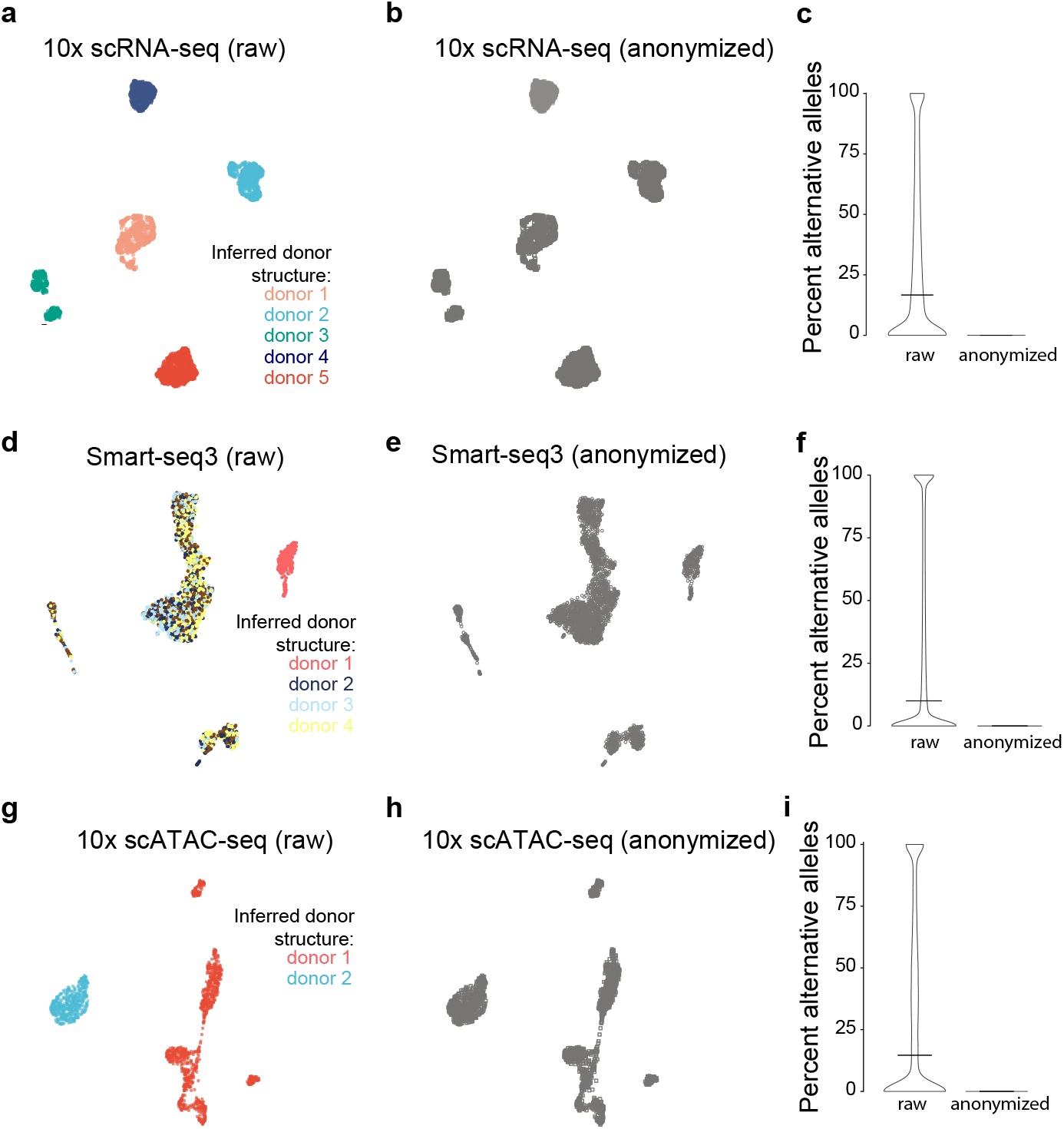
De-identification of human scRNA-seq and scATAC-seq data with *anonymizeBAM*. (**a-b**) UMAP plot of scRNA-seq data (10x Genomics) from five different cell lines (H2228, H1975, A549, H838 and HCC827; in total 2,937 cells), colored according to donor identity inferred from the (a) raw and (b) anonymized sequencing data. (**c**) Percent of 10x Genomics scRNA-seq reads containing alternate allele information at interrogated variant positions, for raw and anonymized data, respectively. All cells were combined and variants with sufficient coverage (n=75,461) were used. (**d-e**) UMAP plot of scRNA-seq data (Smart-seq3) generated from PBMCs (four donors) and HEK293T cells (in total 3,129 cells), colored according to donor identity inferred from the (d) raw and (e) anonymized sequencing data. (**f**) Percent of Smart-seq3 scRNA-seq reads containing alternate allele information at interrogated variant positions, for raw and anonymized data, respectively. All cells were combined and variants with sufficient coverage (n=1,503,063) were used. (**g-h**) UMAP plot of scATAC-seq data (10x Genomics) generated from one PBMC donor and GM12787 cells (in total 1,497 cells), colored according to donor identity inferred from the (d) raw and (e) anonymized sequencing data. (**i**) Percent of 10x Genomics scATAC-seq reads containing alternate allele information at interrogated variant positions, for raw and anonymized data, respectively. All cells were combined and variants with sufficient coverage (n=95,127) were used.

Similar results were achieved when analyzing scRNA-seq data from human peripheral mononuclear cells (PBMCs) of four donors mixed with a HEK293T cell line generated using the Smart-seq3 protocol^19^ (**Figure 2d-f**). To demonstrate that *anonymizeBAM* can also correctly de-identify other data types, we next analyzed single-cell ATAC-seq (scATAC-seq) data that was derived from a single donor of PBMCs and a lymphoblastoid cell line. The donor structure that is clearly present in the raw BAM file (**Figure 2g**) was lost upon anonymization (**Figure 2h**) after which no allele-informative reads remained (**Figure 2i**).

It is crucial that tools used for the de-identification of genomic information work accurately in order to be trusted to be applied for open data sharing.

The strategy we implemented correctly handles SNPs, indels, clipping and multimapped secondary alignments for spliced and regular alignments. We naturally did not attempt to remove copy-number variations (CNVs) in the *anonymizeBAM* tool, since this information is also present in the count files that are currently shared openly.

We here provide the straightforward generation of anonymized sequencing read data with *anonymizeBAM*, which will again enable wide-spread open sharing of sequencing data. This will be important for large publicly funded landmark projects, such as the Human Cell Atlas^20^. Ideally, controlled-access databases (e.g. dbGaP^6^ or EGA^7^) could also directly offer free download of anonymized data as a complement to the full genetic sequence data under controlled access. Finally, our versatile strategy for reliable sequence data anonymization is implemented into a user-friendly and freely available tool present on GitHub (https://github.com/sandberg-lab/dataprivacy) and available from the PyPI repository (can be installed with: *pip install anonymizeBAM*).

## Author contribution

Conceived the idea: R.S. and C.Z. Designed and developed *anonymizeBAM*: C.Z. Performed analyses and generated figures: C.Z. Wrote the manuscript: C.Z and R.S.

## Code availability statement

The code has been deposited on GitHub (https://github.com/sandberg-lab/dataprivacy) and is available on the PyPI repository.

## Data availability statement

No new data was generated in this study.

## Acknowledgements

We would like to thank the members of the Sandberg lab for fruitful discussions. This work was supported by grants to R.S. from the Swedish Research Council (2017-01062), the Knut and Alice Wallenberg Foundation (2017.0110), the Göran Gustafsson Foundation, the National Institute of Health (R01MH109556), and the Bert L. and N. Kuggie Vallee Foundation.

## Code availability

The anonymization procedure has been implemented into a python program and has been deposited in GitHub (https://github.com/sandberg-lab/dataprivacy) and PyPI (https://pypi.org/project/anonymizeBAM).

## Methods

### Data sources

Raw data in fastq format for the 10x Genomics v2 single-cell RNA sequencing experiment of five human cell lines (H2228, H1975, A549, H838 and HCC827) was obtained from the European Nucleotide Archive (accession: SRR8606521). Raw fastq data from scRNA-seq data generated using the Smart-seq3 protocol from PBMC and HEK cells were obtained from ArrayExpress (accession: E-MTAB-8735). Single-cell ATAC-seq data for human PBMCs and GM12878 cells was downloaded as preprocessed (CellRangerATAC v 1.2.0) .bam files from the 10x Genomics website (https://support.10xgenomics.com/single-cell-atac/datasets/1.2.0/atac_hgmm_1k_nextgem; https://support.10xgenomics.com/single-cell-atac/datasets/1.2.0/atac_pbmc_1k_nextgem) and concatenated for the remainder of the analysis.

### scRNA-seq data processing

Raw fastq files were processed using zUMIs^16^ (v 2.9.4f), which relied on mapping of the reads using STAR^11^ (v 2.5.4b) to the human genome (hg38) and quantification using Ensembl gene annotations (GRCh38.95). To generate UMAP plots, count tables were analyzed in Seurat^21^ (v.3.1.5) following the standard workflow with default settings. In the case of Smart-seq3 data, the UMAP coordinates as previously published were used^19^.

### anonymizeBAM workflow

The anonymization procedure involves modification of the observed read sequence to the reference genome sequence and de-identification of auxiliary tags.

Overview of considered cases and the associated anonymization strategy:

1. SNPs: Mismatches to the reference (either explicitly *X* coded in the CIGAR value or within *M* matched segments) are replaced by the reference base.
2. Insertions (CIGAR *I*): The read sequence is extended by the length equal to the insertion, while keeping the 5’ mapping position constant.
3. Deletions (CIGAR *D*): The missing reference sequence is inserted into the read, while removing an equal number of bases from the 3’ end.
4. Clipping: soft or hard clipped bases (CIGAR: *S* / *H*) are replaced by reference sequence of matching length. If reads start with clipped bases in single-end data, the reference position of the read start is adjusted, which is not possible for paired-end reads to conserve correct matepair information in the TLEN and PNEXT fields. Thus, for paired-end reads, the clipped sequence length is added at the end of the read.
5. Splicing: Splicing is observed and splice-sites are conserved even in the case of deletions and insertions. In case of a deletion leading to a shift of length that is longer than the mapped sequence length in the last exon, this splice event is removed.
6. Multimapping: In default behavior, only primary alignments are emitted. The user can choose to keep secondary but we note that full anonymization cannot be guaranteed.
7. Unmapped reads: Unmapped reads cannot be sanitized and are discarded in default settings.

As donor-related information could also be inferred from standard bam fields and auxiliary tags, the following changes are made:

1. CIGAR value is matched to the anonymized sequence (Example: 100M).
2. MD tags coding mismatches or deleted information are updated to the anonymized sequence, if present (Example: 100).
3. NM and nM tags (edit distance to the reference) are cleared by replacement with 0.
4. Tags containing information on the alignment are discarded (MC, XN, XM, XO, XG). In *--strict* mode, the following tags are also changed:
5. Mapping quality is set to maximum/unavailable (255).
6. AS and MQ (alignment score / mapping quality) are set to read length.
7. NH (number of hits in the reference) is set to 1.
8. Discarding of the following tags: HI, IH, H1, H2, OA, OC, OP, OQ, SA, SM, XA, XS.

### Genotyping and donor inference

Genotype-informative base coverage was summarized per cell over 7.4 million common variants (AF>5% in the 1000 genomes project phase 3) with cellsnp-lite^17^ (v 1.2.0) using the --UMItag None --minCOUNT 10 settings. The resulting sparse VCF file was loaded into vireo^18^ (v 0.4.2) and donor deconvolution performed using the appropriate --nDonor flag (n = 2 for scATAC, n=5 for 10x scRNAseq and n=6 for Smart-seq3).

### Validation BAM specification

The output of *anonymizeBAM* from the Smart-seq3 single-cell RNA-seq dataset was validated for compliance with the SAM specification^13^ using the picard-tools (http://broadinstitute.github.io/picard) ValidateSamFile command with the following exceptions: --IGNORE MATE_NOT_FOUND, RECORD_MISSING_READ_GROUP, MISSING_READ_GROUP. No errors or warnings were observed.

## References

1. Shendure, J. et al. DNA sequencing at 40: past, present and future. Nature 550, 345–353 (2017).

2. Stark, R., Grzelak, M. & Hadfield, J. RNA sequencing: the teenage years. Nat. Rev. Genet. 20, 631–656 (2019).

3. Svensson, V., da Veiga Beltrame, E. & Pachter, L. A curated database reveals trends in single-cell transcriptomics. Database 2020, (2020).

4. Erlich, Y., Shor, T., Pe’er, I. & Carmi, S. Identity inference of genomic data using long-range familial searches. Science vol. 362 690–694 (2018).

5. Shabani, M. & Marelli, L. Re-identifiability of genomic data and the GDPR: Assessing the re-identifiability of genomic data in light of the EU General Data Protection Regulation: Assessing the re-identifiability of genomic data in light of the EU General Data Protection Regulation. EMBO Rep. 20, e48316 (2019).

6. Tryka, K. A. et al. NCBI’s Database of Genotypes and Phenotypes: dbGaP. Nucleic Acids Res. 42, D975–9 (2014).

7. Lappalainen, I. et al. The European Genome-phenome Archive of human data consented for biomedical research. Nat. Genet. 47, 692–695 (2015).

8. Ozercan, H. I., Ileri, A. M., Ayday, E. & Alkan, C. Realizing the potential of blockchain technologies in genomics. Genome Res. 28, 1255–1263 (2018).

9. Joly, Y., Dyke, S. O. M., Knoppers, B. M. & Pastinen, T. Are Data Sharing and Privacy Protection Mutually Exclusive? Cell 167, 1150–1154 (2016).

10. Gürsoy, G. et al. Data Sanitization to Reduce Private Information Leakage from Functional Genomics. Cell 183, 905–917.e16 (2020).

11. Dobin, A. et al. STAR: ultrafast universal RNA-seq aligner. Bioinformatics 29, 15–21 (2013).

12. Li, H. & Durbin, R. Fast and accurate short read alignment with Burrows-Wheeler transform. Bioinformatics 25, 1754–1760 (2009).

13. The SAM/BAM Format Specification Working Group. Sequence alignment/map format specification. http://samtools.github.io/hts-specs/SAMv1.pdf.

14. Tian, L. et al. Benchmarking single cell RNA-sequencing analysis pipelines using mixture control experiments. Nat. Methods 16, 479–487 (2019).

15. Zheng, G. X. Y. et al. Massively parallel digital transcriptional profiling of single cells. Nat. Commun. 8, 14049 (2017).

16. Parekh, S., Ziegenhain, C., Vieth, B., Enard, W. & Hellmann, I. zUMIs - A fast and flexible pipeline to process RNA sequencing data with UMIs. Gigascience 7, (2018).

17. Huang, X. & Huang, Y. Cellsnp-lite: an efficient tool for genotyping single cells. Cold Spring Harbor Laboratory 2020.12.31.424913 (2021) doi:10.1101/2020.12.31.424913.

18. Huang, Y., McCarthy, D. J. & Stegle, O. Vireo: Bayesian demultiplexing of pooled singlecell RNA-seq data without genotype reference. bioRxiv (2019).

19. Hagemann-Jensen, M. et al. Single-cell RNA counting at allele and isoform resolution using Smart-seq3. Nat. Biotechnol. (2020) doi:10.1038/s41587-020-0497-0.

20. Regev, A. et al. The Human Cell Atlas. Elife 6, (2017).

21. Stuart, T. et al. Comprehensive integration of single-cell data. Cell 177, 1888–1902.e21 (2019).

